# BPscore: an effective metric for meaningful comparisons of structural chromosome segmentations

**DOI:** 10.1101/307330

**Authors:** Rafal Zaborowski, Bartek Wilczyński

## Abstract

Studying the 3D structure of chromosomes is an emerging field flourishing in recent years because of rapid development of experimental approaches for studying chromosomal contacts. This has led to numerous studies providing results of segmentation of chromosome sequences of different species into so called Topologically Associating Domains (TADs). As the number of such studies grows steadily and many of them make claims about the perceived differences between TAD structures observed in different conditions, there is a growing need for good measures of similarity (or dissimilarity) between such segmentations. We provide here a BP score, which is a relatively simple distance metric based on the bipartite matching between two segmentations. In this paper, we provide the rationale behind choosing specifically this function and show its results on several different datasets, both simulated and experimental. We show that not only the BP score is a proper metric satisfying the triangle inequality, but that it is providing good granularity of scores for typical situations occuring between different TAD segmentations. We also introduce local variant of the BP metric and show that in actual comparisons between experimental datasets, the local BP score is correlating with the observed changes in gene expression and genome methylation. In summary, we consider the BP score a good foundation for analysing the dynamics of chromosome structures. The methodology we present in this work could be used by many researchers in their ongoing analyses making it a popular and useful tool.

**Author summary:** Many researchers are interested in the chromosomal structure, its function and dynamics. Over the recent years, chromosome conformation capture (3C) methods have become the main source of experimental data on the subject and the Topologically Associating Domains (TADs) have become the *de-facto* standard unit of chromosomal structure. Many methods have been developed for TAD calling and the 3C experiments have been done in multiple conditions giving us a multitude of chromosomal segmentations describing the most atomic differences in chromosomal structure between conditions. Until now, such segmentations were compared mostly by very rough measures, such as the Jaccard coefficient or TAD overlaps or very general metrics like the variation of information coefficient. This has limited the researchers in the analysis of differential TAD segmentations, and practically prevented any proper analysis of TAD dynamics between conditions. Our approach has the potential to facilitate such analyses by providing researchers with mathematically sound metric that is designed specifically for the purpose and tested on both simulated and experimental data. Additionally, we provide a local variant of our measure that is a natural derivative of the BP score that can indicate which parts of the chromosomes are undergoing the most significant structural reorganizations.

## Introduction

Scientific studies of the cell nucleus and its structure started more than a hundered years ago when embryologists were performing their experiments on the fertilized eggs and subsequent rapid cell divisions. In the last 50 years, after discovery of the DNA structrue and its role, the topic of chromatin structure was considered to be of slightly lower importance, as the researchers have been mostly preoccupied with studying the DNA sequence of the genomes to uncover its function. In recent years, genome conformation studies gained significant attention again, for at least two reasons. On the technical side, this is mainly due to development of Chromosome Conformation Capture, or 3C technique [1] and its derivative methods like Hi-C [2], 4C [3,4] or 5C [5]. On the other hand, a perhaps more important reason why there is so much interest in the newly acquired data on chromosomal conformation is that current efforts in explaining the function of non-coding regulatory elements without the information on the contacts between enhancers and promoters are proving it to be an extremely difficult problem [19]. The problem of gene regulation is difficult even given that the positions of such elements can be identified by DNAse Seq or related techniques [20]. This leads to the situation where many researchers are probing chromosome structure using 3C related techniques and the research in the field is progressing rapidly. This, as always is the case in blooming scientific fields, leads to new technical challenges in describing the chromosomal structures and comparing them between cell-types and experimental conditions.

Going back to the technical side of the chromosome capture techniques, in the Hi-C assay DNA is cross-linked, shared into pairs of fragments and ligated. Ligated fragments are then pulled down and merged with adapters. Finally, fragments are paired-end sequenced to produce library of paired reads indicating pairs of genomic regions that are in close proximity in the three-dimensional space of the nucleus. Even though the method is not exact in the sense of mapping the 3D-distance to the probability of ligation followed by succesful identification of a pair, it has been proven to be very reproducible provided that we are interested in the average contact frequency over a large pool of cells statistically sampled by deep sequencing [21]. Single cell hi-c techniques are much more recently developed, but there is significant progress there as well and we should expect soon to be able to compare chromosomal contact structures also between individual cells [22] [23]. Also, other methods, that are not ligation-dependent have been developed for probing chromosomal contacts, most notably the genome architecture mapping (GAM) results corroborate the hi-c derived chromosomal structures [24].

Once the sequencing is completed and contacting pairs of segments are identified, usually the analysis of interactions is based on construction of contact maps: symmetric matrices that summarize contacts between bins of a pre-defined size. If we are interested in cis-chromosomal contacts, which is usually the case, such large genome-wide matrix is usually broken down into separate matrices for each chromosome. In these matrices, each cell contains the number of interactions between 2 particular regions of the same chromosome. Such matrices usually need to be normalized, to account for several technical issues, such as GC content bias and diverse mappability across genomic regions that lead to non-uniform read coverage along chromosomes. There are methods that are specialized to a given hi-c protocol [25] as well as more generic normalization methods based on the assumption of uniform coverage [16].

In recent years, many studies have suggested that in addition to the well established chromosomal compartments representing the division of chromatin into active euchromatin and inactive heterochromatin, there exists a finer structure of Topologically Associating domains or TADs for short [6] [26]. The presence of TAD-like structures was confirmed in almost all metazoan Hi-C data, with notable exclusion of some chromosomes in C. elegans [27], while there appear to be no sign of TADs in yeast and plants [28]. The majority of animals, including all studied mammals, show significant concentrations of contacts into TADs and these appear to be largely conserved between species and conditions [6]. Even though, mostly due to variablity in hi-c protocols and TAD-calling methods, it still remains a challenge to assess physical properties and biological function of TADs in individual cells, it was shown beyond doubt that regions of genome demarcated by TADs are indeed correlated with gene regulation [7]. In particular it was shown that a deletion of TAD boundary regions may lead to different developmental disorders [8] [9].

Numerous studies exploiting Hi-C to compare genome conformation between different cell types or conditions were conducted [10–12,15]. The purpose of such analyses is to capture differences in spatial organization of chromatin and its relationships with gene expression or epigenetic modifications. One approach is to spot and quantify regions of chromosome where TAD arrangement is (dis)similar between two experiments. A common method for this purpose is to count number or overlapping TAD boundaries that was used among others by [6] [13] [14]. However this approach suffers one serious disadvantage that large domains with relatively small shift in their boundaries contribute to lowering the overall dissimilarity as much as small domains with relatively large shifts. Another approach, that is not as boundary-centric is by measuring variation of information between different TAD-segmentations as used in [15]. This approach, however suffers from the fact that it does not take into account the linear structure of the genome and usually overestimates the deviation from the randomized control and therefore overestimates the TAD structure conservation. Given the rapid advancement of the experimental research in the field and constantly growing flow of new data with TAD structures, it is important to have a proper metric to compare different experimental results as well as different computational methods for TAD detection. In perfect scenario, such dissimilarity measure should be a proper metric satisfying triangle inequality.

In this work we present new distance measure for comparing TAD sets and prove that our measure fulfill metric properties. We use simulated and real data to show that our metric is able to capture interesting properties of different domain sets. We also perform comparison with simple boundaries overlap count based approach concluding that our metric exhibit smoother behaviour allowing for more precise analysis. Finally we present how our metric can be used to estimate (dis)similarity of subchromosomal regions enabling for more in depth analysis.

## Results

### BP-distance

In this chapter we give the definition of BP distance and explain notation. Precise definitions can be found in chapter Materials and methods.

Let *A* and *B* define 2 intervals of equal length. Each of them is partitioned on subintervals *A*[*i*] and *B*[*j*] (see fig. 1). A pair of subintervals *A*[*i*], *B*[*j*] may induce segment *o*[*k*], which is non empty intersection between those subintervals. BP distance between domain sets *A* and *B* is defined as:

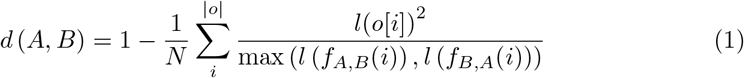

where *f_A,B_*(*i*) is a function mapping *o*[*i*] to subinterval in *A*, which induced *o*[*i*] and *l*(*o*[*i*]) is function mapping *o*[*i*] to its length.

**Fig 1.**
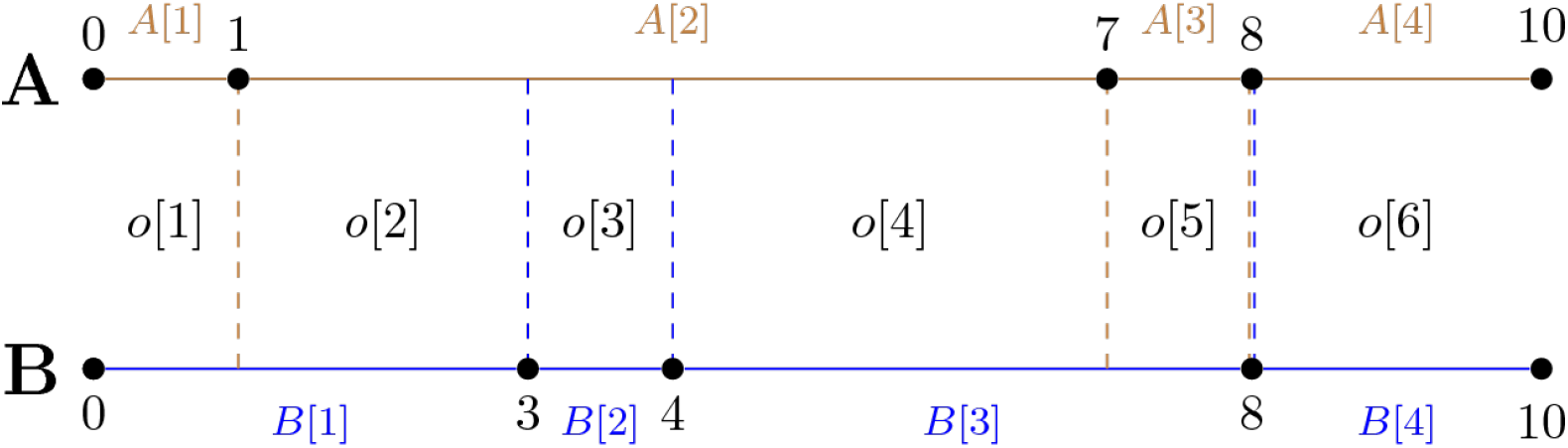
Chromosome partitioning and the induced segmentation.

**Fact 1.** *The above defined function is a metric*.

The proof can be found in chapter Materials and methods. Below we present some properties of function *d*(*A, B*):

1. every change that is introduced has proportional impact to the length of the interval affected,
2. inclusions, deletions and shifts are treated equally,
3. we can compare domain sets of different sizes.

The above properties help avoid problems, which arise in alternative TADs similarity measures, where one assess similarity based on the number of TAD boundary overlaps across chromosome (see next section).

### Comparison with Existing Methods

To demonstrate advantage of our metric over boundaries overlap count approach we simulated artificial sets of TADs using following procedure. First we simulate 2 sets, each consisting of 600 TADs, the approximate number of TADs on chromosome 1 determined using DP approach [15] on human ESC Hi-C data [6]:

1. The TADs lengths were drawn to reflect real TAD length distribution (Negative Binomial distribution with parameters adjusted manually),
2. TAD length was fixed to 40 bins for every TAD.

Then for each of the above 2 sets we generated match sets according to 3 scenarios:

- each domain boundary is moved at most *ε* bins left or right of its initial position (fig. 2a),
- new boundary is introduced at most *ε* bins to the left of each TAD boundary, except of first one (fig. 2b),
- 2^*n*^ – 1 boundaries are introduced into every TAD according to binary interval partitioning scheme (fig. 2c).

Moreover each of the above match sets is replicated 3 times with increasing parameter *ε* or *n*. Scenario 1 can represent situation where structural organization of 2 chromosomes is almost identical however due to noise factors coordinates of detected TADs do not overlap. Scenario 2 can be considered either a noisy match (just like 1) or large mismatch. The former case may occur when a little interval like non-TAD region (often produced in Directionality Index method [6] or DP approach [15]) is located next to large domain and both of them overlap with some large domain in corresponding TAD set. The latter one is possible when 2 consecutive TADs of similar length overlap with 1 large TAD, a situation indicating poor match. Scenario 3 can be considered mismatch except for *n* = 0. In particular we can imagine complete mismatch if a TAD in experiment A have length *N* and in experiment B there are *N* TADs, each of length 1.

**Fig 2.**
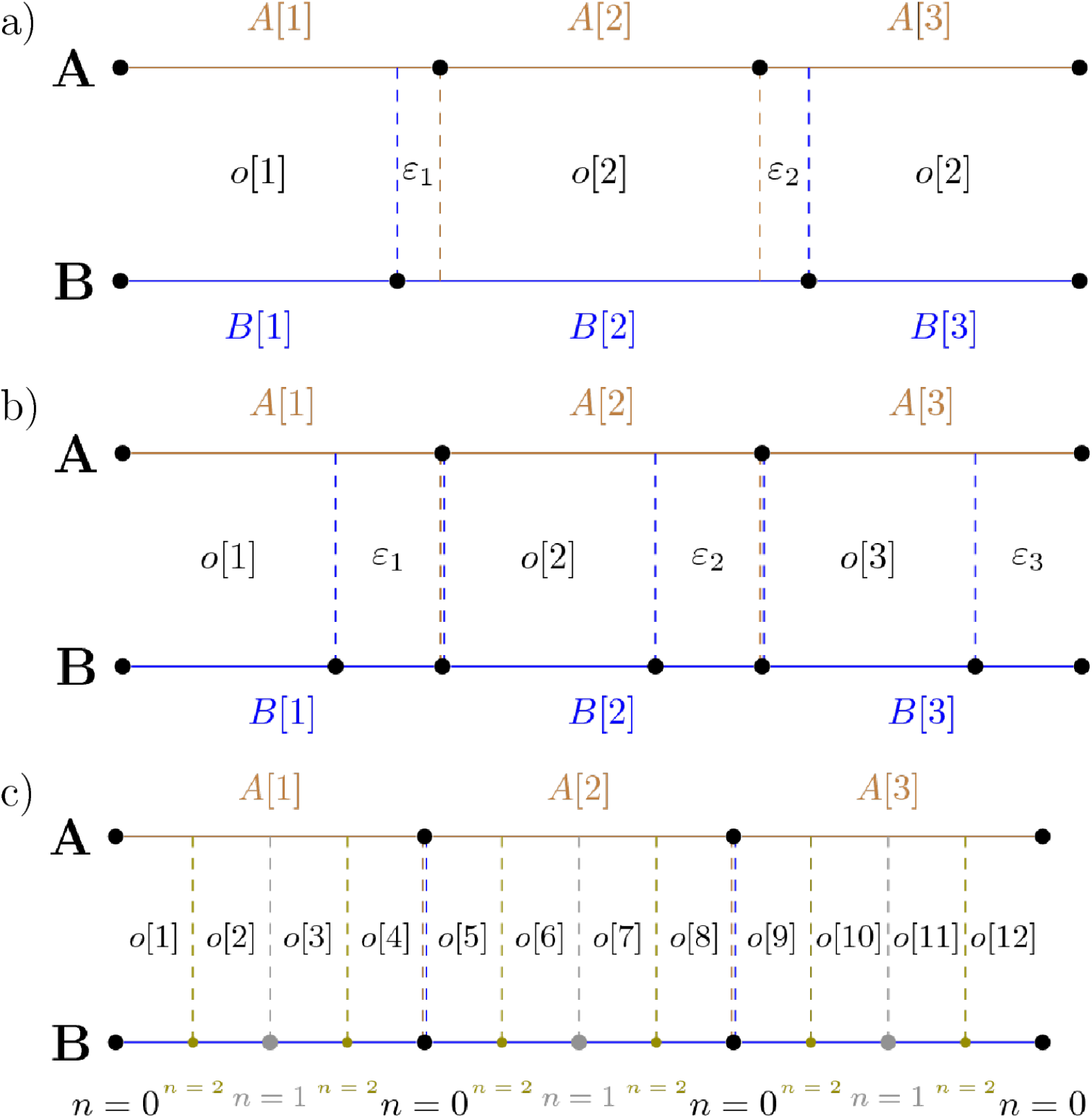
*ε*-matching scenarios. **a)** Boundaries in *B* are moved at most *ε* bins left or right. **b)** New boundary is inserted into *B* at most *ε* bins to the left of existing boundary. **c)** Domains in *B* are partitioned according to binary interval partitioning scheme.

Additionally we compare our metric with Variation of Information metric adopted to assess similarity of segmentation as described in [15].

For each scenario and possible parameter value we report 3 numbers (fig. 3): our distance (refered to as BP), Jaccard Index distance (refered to as JI) and Variation of Information distance (refered to as VI). The JI distance is calculated by dividing the number of TAD boundary overlaps between *A* and *B* by the number of all boundaries (overlaps are counted once) and subtracting resulting fraction from one. Calculation of VI distance is described in Materials and methods.

**Fig 3.**
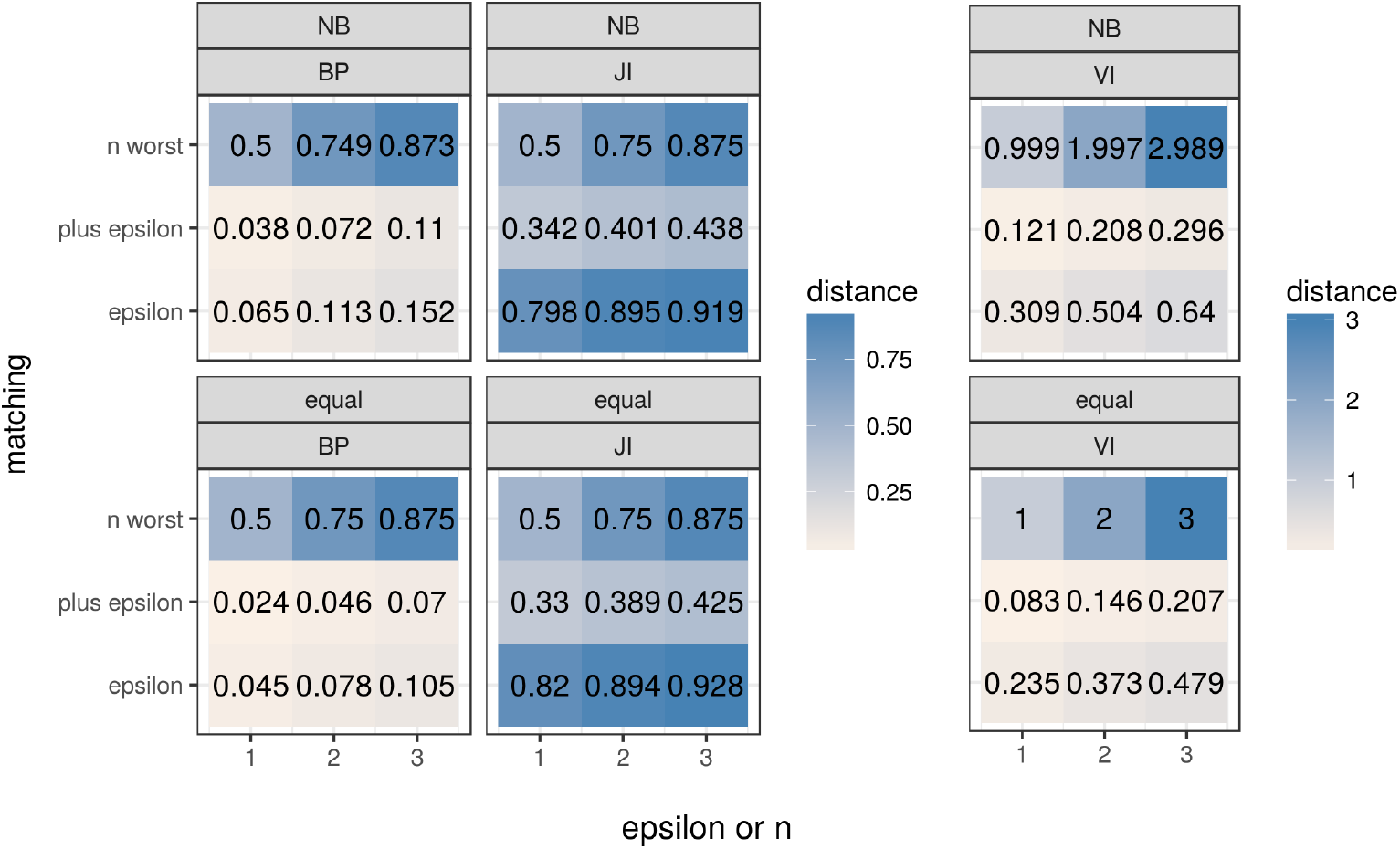
Comparison with Jaccard Index and Variation of Information distance.

As can be seen in situations where perfect TADs overlap is disturbed by subtle noise the BPscore distance, similarly to the VI distance, outperforms boundary overlap count (Jaccard Index) approach in recovering true similarity.

### Differentiating Technical From Biological Replicates Data

To demonstrate usefulness of our metric we present a case where one might be interested in assessing structural similarities genomewide for multiple samples. We selected publicly available sets of Hi-C data consisting of 6 cell lines: ESC, MSC, MDC, NPC, TDC and IMR90 [6] [11]. ESC was available in 4 technical replicates and remaining lines in 2 technical replicates. We processed each dataset using Armatus software [15] to produce sets of TADs. Finally we made pairwise comparisons for every pair of TAD sets. Figure 4 illustrates comparisons divided on two categories: within replicates and between cells for BP metric (left), JI metric (center) and VI metric (right). As can be seen all metrics discriminate between two groups under consideration, i. e. the scores of between-replicates comparisons are consistently lower than between-conditions comparisons. While all metrics give significant discrimation, we note that the significance of the difference, as well as the separation between the two groups is better for the VI and BP scores than the Jaccard index. Of the two metrics with better separation, BP score is giving slightly lower p-value, however this is unlikely to be a difference of practical importance. Similar behaviour is observed when comparing individual chromosomes (S1 Fig and S2 Fig).

**Fig 4.**
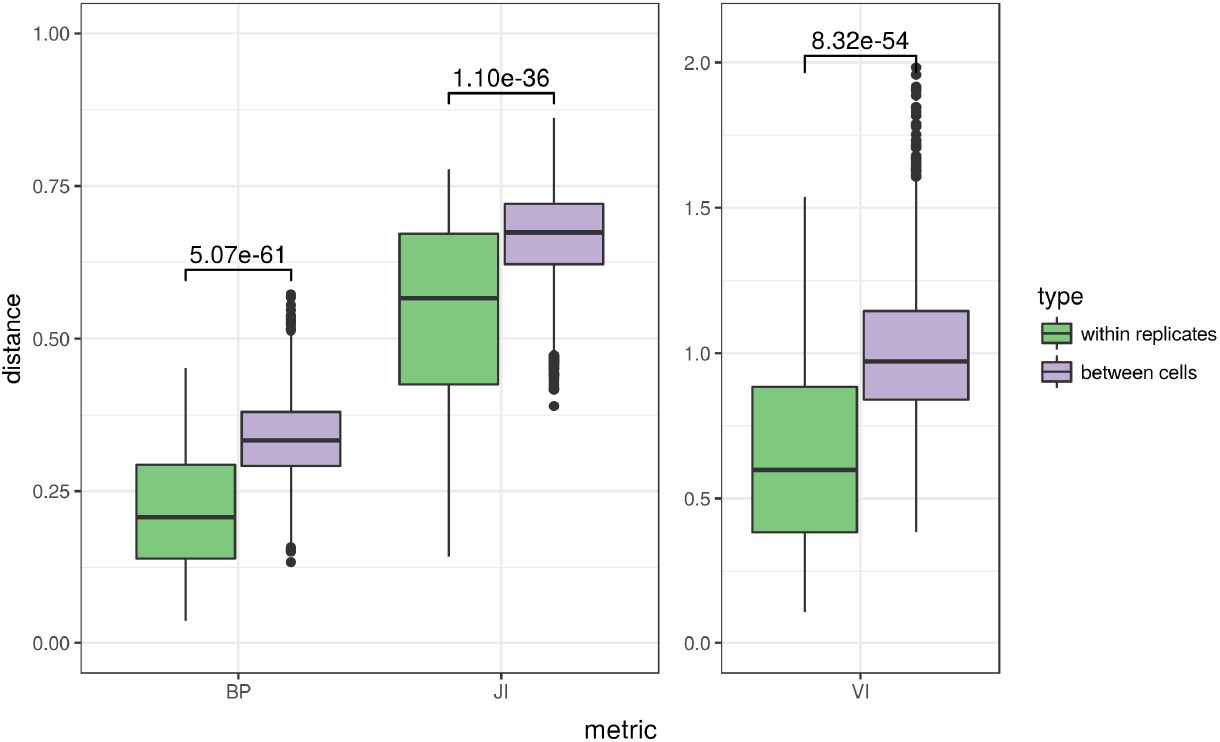
Pairwise comparisons of real Hi-C datasets with different metrics.

### Local BP score

Another potentially important application of comparing chromosomal segmentations is the search for locally re-organized fragments induced by 2 partitions for selected fragment of a chromosome. For example to check if differential gene expression or methylation pattern correlates with segmentation pattern. For this kind of analysis we suggest to use local BP score - a measure associated with our metric. Local BP score of segment *i* is defined as:

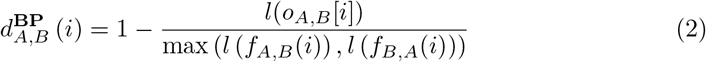

Local BP score satisfies: 0 ≤ *d*_*A,B*_ (*i*) < 1. The closer it is to 0, the larger the overlap between *f_A,B_*(*i*) and *f*_*B*,*A*_(*i*). Conversely, the closer it is to 1, the higher the mismatch. Value of 0.5 may represent a boundary situation between match and mismatch case. Fig. 5a illustrates segmentation of human chromosome 1 (100 to 321 bins fragment in 40 kb resolution) between ESC and MSC cells. Bottom axis ticks represent bins, top triangles are TAD boundaries from MSC and bottom triangles are TAD boundaries belonging to ESC. Black, dashed vertical lines mark partitions, which increase matching (TAD boundaries in both sets overlap) while grey, dashed vertical lines denote partitions increasing mismatch.

**Fig 5.**
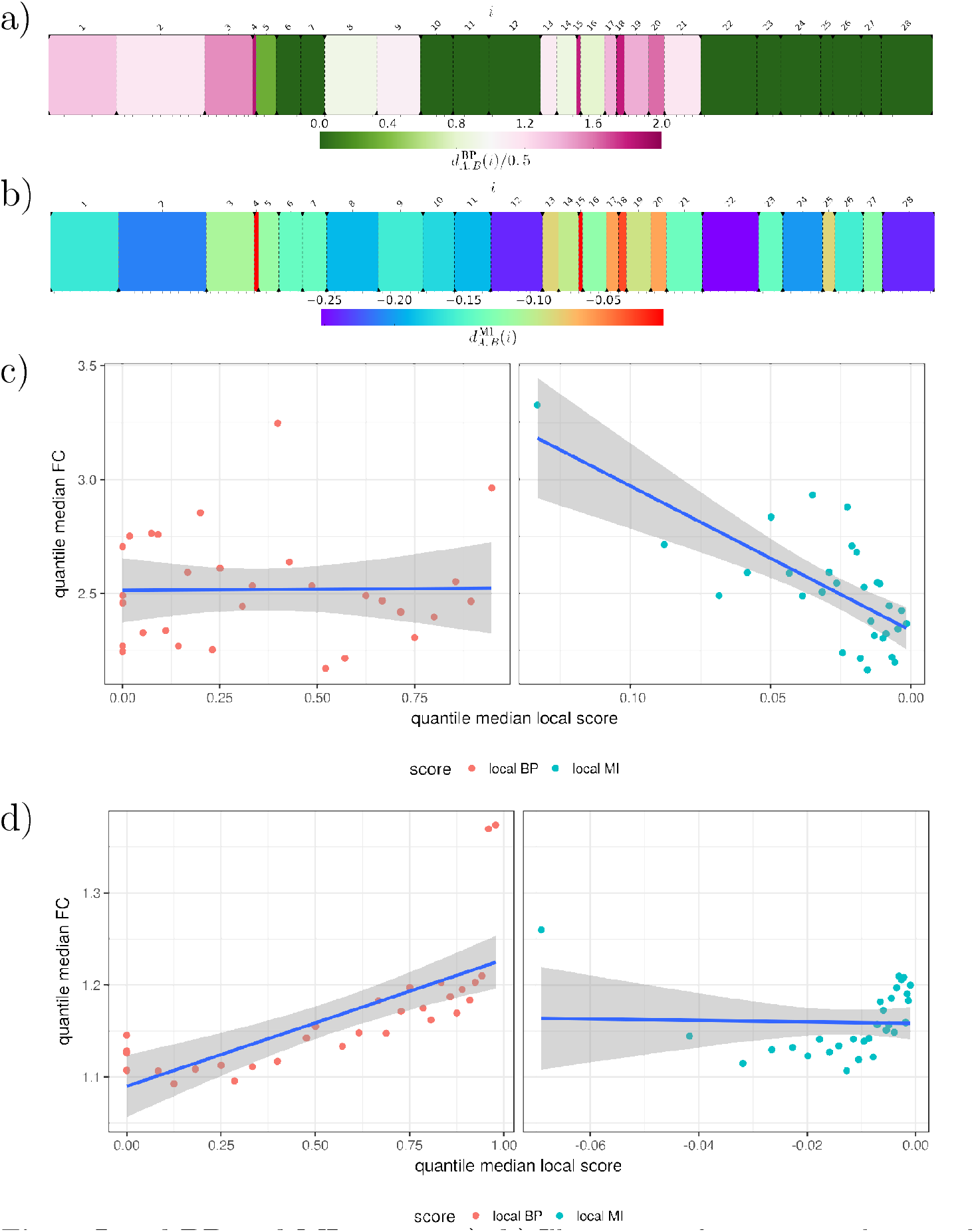
Local BP and MI scores. **a), b)** Illustration of partitioning between human ESC and MSC TADs of 4000 kb - 12840 kb (40 kb resolution) region of chromosome 1 with local BP score and local MI score respectively. **c)** Gene expression fold change vs local BP score or local MI score of 30 quantiles between human IMR90 and NPC cells. **d)** Same as in c), but with methylation fold change instead of gene expression.

Although in [15] the authors only consider the VI measure for global segmentation comparisons and do not discuss any local version of this measure, we thought that a natural measure of local segmentation related to Variation of Information would be local MI (Mutual Information) expressed with following formula:

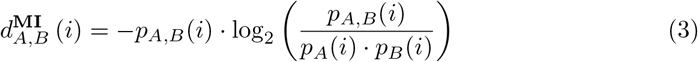

where: 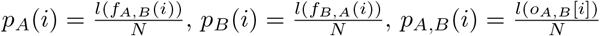. In local MI score segments tend to be first ordered by their size (descending) and second by similarity of domains, which induce them (from match to mismatch), see fig. 5b.

Based on hypothesis that TADs define regulatory landscapes and may limit activity of certain regulatory elements [8] [9] we decided to check if there is a correlation between local score and differential gene expression or methylation pattern genome wide. To validate our prediction we derived differential gene expression and methylation data from publicly available resources. We then created all by all pairing of hi-c datasets for available cells and for each pair we assigned local BP and MI scores and fold change of gene expression or methylation for every gene (S3 Fig, S8 Fig). Due to large amount of genes we aggregated genes into 30 quantiles based on the local score value to reduce noise. Then for each group we plotted its quantile versus mean fold change (with standard deviation). This revealed corelations between gene expression or methylation fold change and the local domain rearrangement score (S4 Fig, S9 Fig).

Finally fig. 5c and fig. 5d illustrates the relationship between fold change of gene expression and methylation respectively as a function of local score using IMR90 and NPC cells as the example. Here genes were grouped based on local score so in summary there are 30 groups (quantiles). For each group median local score vs median fold change is shown. As can be seen the relationship is not strong, albeit visible and significant. Images for remaining pairs of cells can be found in supplement (S5 Fig, S10 Fig). The visible tendency of the local BP score to better correlate with methylation fold change and the MI score to better correlate with gene expression is generally representative of most datasets we analyzed. Corelations measured using PCC and Spearmans rho as well as significances are illustrated in S6 Fig and S7 Fig for gene expression, S11 Fig and S12 Fig for methylations and summarized in S1 Table and S2 Table.

## Discussion

In this paper, we have considered the problem of comparisons between different segmentations of chromosomes into TADs. Currently, such comparisons are mainly based on comparing the locations of TAD boundaries and quantification of discrepancies between two sets of such locations. This is usually done using the Jaccard coefficient or some analogous function. We show in this work, that under a number of realistic scenarios, where the differences between TAD segmentations are driven by experimental noise, this is not an effective strategy. By utilising different e perturbations of TAD segmentations, we can show that Jaccard Index is a poor technique for discerning significantly changed segmentations, from those with minor perturbations.

We provide an alternative scoring scheme, so called BP score, deriving its name from the bi-partite segment overlap graph. This score has several advantages over the Jaccard Index. Firstly, it is naturally scaling with the size of chromosomes and the resolution at which the segmentation is done, that is any change in the TAD structure will contribute to the metric proportionally to its size. Additionally, the method is treating equally all kinds of changes (insertions, deletions and shifts of domain boundaries) and can be meaningfully used to compare domain sets with different granularity, i.e. containing significantly different numbers of domains. On the technical level, the BP score is a proper distance metric, which has advantages for applications that need symmetry or triangle inequality, including clustering methods. The global BP metric is similar in the applicability to the Variation of Information metric suggested for the task earlier [15]. Not only are these methods giving more sensible results than JI in the three simulated scenarios of *ϵ*-matchings, but are also giving us a better discernibility between segmentations derived from experimental hi-c replicates and actual segmentations originating from different experimental conditions. This shows, that the researchers trying to obtain a meaningful score for TAD segmentation divergence should have more granular results with BP and VI scores than with JI or similar approaches.

In addition to the global VI and BP scores, we propose the analysis of local versions of the two: local BP and local MI scores. The usefulness these local measures of domain rearrangement is supported by our analysis of experimental data. We show that in actual comparisons between different TAD segmentations performed on different cell lines, the regions with high local BP and local MI scores, i.e. those identified by the BP and MI score as the most rearranged, show the highest fold-changes in gene expression and methylation changes. This might be very useful in uncovering the mechanisms connecting chromosome structure rearrangement to functional changes like gene expression.

We provide an implementation of the BP, VI and JI scores that allows all researchers to test them on their own data. The implementation of the BP score is relatively simple as it can be computed as a sum of local scores for each of the atomic segment induced from the two segmentations being compared. This allows also to identify local contributions to the BP score, therefore leading to the identification of fragments of chromosomes that yield the most divergent parts of the chromosomes between the two segmentations under consideration. This might be specifically useful to the researches studying chromosomal domain dynamics, allowing them to focus their attention on parts of chromosomes that undergo the most dynamic TAD rearrangements.

## Conclusion

All these results together show that the BP score is a valid metric for comparisons of TAD segmentations. It is superior to the widely used Jaccard Index, and can be useful in many comparative studies of chromatin domains. Not only is the BP score a better metric than the JI for the global comparison of segmentations, but its local version can give us an interesting measure of domain rearrangement that we show to be correlated with functional measurements.

We have focused our analyses of the BP score solely on its application to the TAD segmentations originating from hi-c experiments. This is however not an exclusive application of this measure. Mathematically, there is nothing in the BP score that would limit its adoption in the field of other chromosome segmentation situations, such as haplotype inference, epigenetic domains and many others.

## Materials and methods

### Hi-C data and its preprocessing

Raw Hi-C data was downloaded from Gene Expression Omnibus, repositories: GSE35156 and GSE52457. Reads were processed using Iterative Correction [16] pipeline: mapping, bining, filtering, contact maps generation and iterative correction steps with default parameters.

### TADs generation

TAD boundaries were determined using Armatus software [15] (with default parameters) and processed to fill inter TAD gaps with artificial domains. As a result a set of TADs on any chromosome comprised sequence of nonoverlapping, consecutive intervals having total chromosome length the same for each cell type. Unmappable regions (contact map row/column sums to zero) were marked and excluded in all later analyses.

### Calculation of Variation of Information

The VI distance between 2 partitions of the same chromosome is calculated as described in [15]. Briefly given 2 partitions *A* and *B* of length *N*:

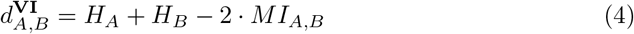

where: *H_A_, H_B_* are entropies of partitions *A* and *B* respectively and *MI_A,B_* is their Mutual Information. Entropy of partition *X* is given by:

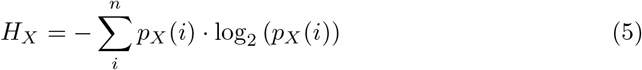

and Mutual Information of partitions *A* and *B* is expressed with:

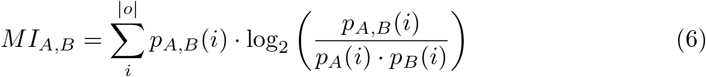

where: 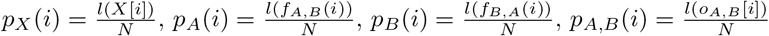 and *n* is a number of domains in *X*.

### Differential gene expression

Gene expression data (bam files) was taken from http://epigenome.ucsd.edu/differentiation/download.html as pointed in [17]. Bam files were processed using edgeR [18] as described in manual to produce differential gene expression.

### Methylation data

Methylations data was taken from: http://epigenome.ucsd.edu/differentiation/download.html [17] and processed using in house scripts to obtain fold changes for each segment in every cell type. Briefly each methylated position was asigned to segment and then for each segment a ratio of methylated by unmethylated counts was calculated.

### Source code

The source coude is publicly available on github: https://github.com/rz6/bp-metric.

### Proof of fact 1

We start with defining concepts used throughout this proof.

**Definition 1.** *A segment is semi-closed discrete interval:* (*a, b*] = {*x* ∈ ℕ^+^ |*a* < *x* ≤ *b*} *with following standard relations*:

i. *equality:* (*a, b*] = (*c, d*] ⟺ *a* = *c* ∧ *b* = *d*
ii. *subset:* (*a, b*] ⊆ (*c, d*] ⟺ *a* ≥ *c* ∧ *b* ≤ *d*
iii. *intersection:*

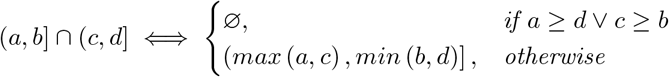

In this study we use segments to describe chromosomes, TADs and overlaps between them.

**Remark 1.** *Each chromosome can be partitioned on collection of non-overlapping, consecutive segments (domains). We restrict notation to one chromosome for simplicity and from now on assume that any partition refer to the same chromosome. This notation can be naturally extended to multiple chromosomes for example by assumming segmentations of concatenated chromosomes with fixed boundaries between them. We will use capital letters to distinguish partitions, and we will use indexes to refer to sorted segments (domains): X* [*i*], *Y* [*j*].

**Definition 2.** *We define following functions on segments:*

i. *segment start: s* ((*a, b*]) = *a*,
ii. *segment end: e* ((*a, b*]) = *b*,
iii. *segment length: l* ((*a, b*]) = *b* – *a*,

As partitions being compared *X, Y*, … refer to the same chromosome we can write *l*(*X*) = *l*(*Y*) = … = *N*.

**Definition 3.** *Intersecting two partitions X and Y induces a partition we call the segmentation o_X,Y_ with following properties:*

i. ∀_*i*_∀_*j*_ *X* [*i*] ⋂ *Y* [*j*] ≠ ∅ ⇒ *X* [*i*] ⋂ *Y* [*j*] ∈ *o*_*X,Y*_
ii. ∀_*i*_∀_*j*>*i*_ *s* (*o*_*X,Y*_ [*i*]) < *s* (*o*_*X,Y*_ [*j*])

**Remark 2.** *When considering triplet partitions X,Y,Z we distinguish between 2 types of segments, atomic and non-atomic (or divisible). We call a segment* (*a, b*] *atomic and write o*[*i*] *if there is no other segment o_X,Y_* [*k*], *o*_*X,Z*_ [*l*] *or o*_*Y,Z*_ [*m*] *that is smaller than* (*a, b*] *and included in* (*a, b*]. *Otherwise we call segment non-atomic and write o_X,Y_* [*i*].

For example in figure 6 segment *o_A,C_* [3] is not atomic - it can be further partitioned into *o*[3] and *o*[4] both of which are atomic.

**Fig 6.**
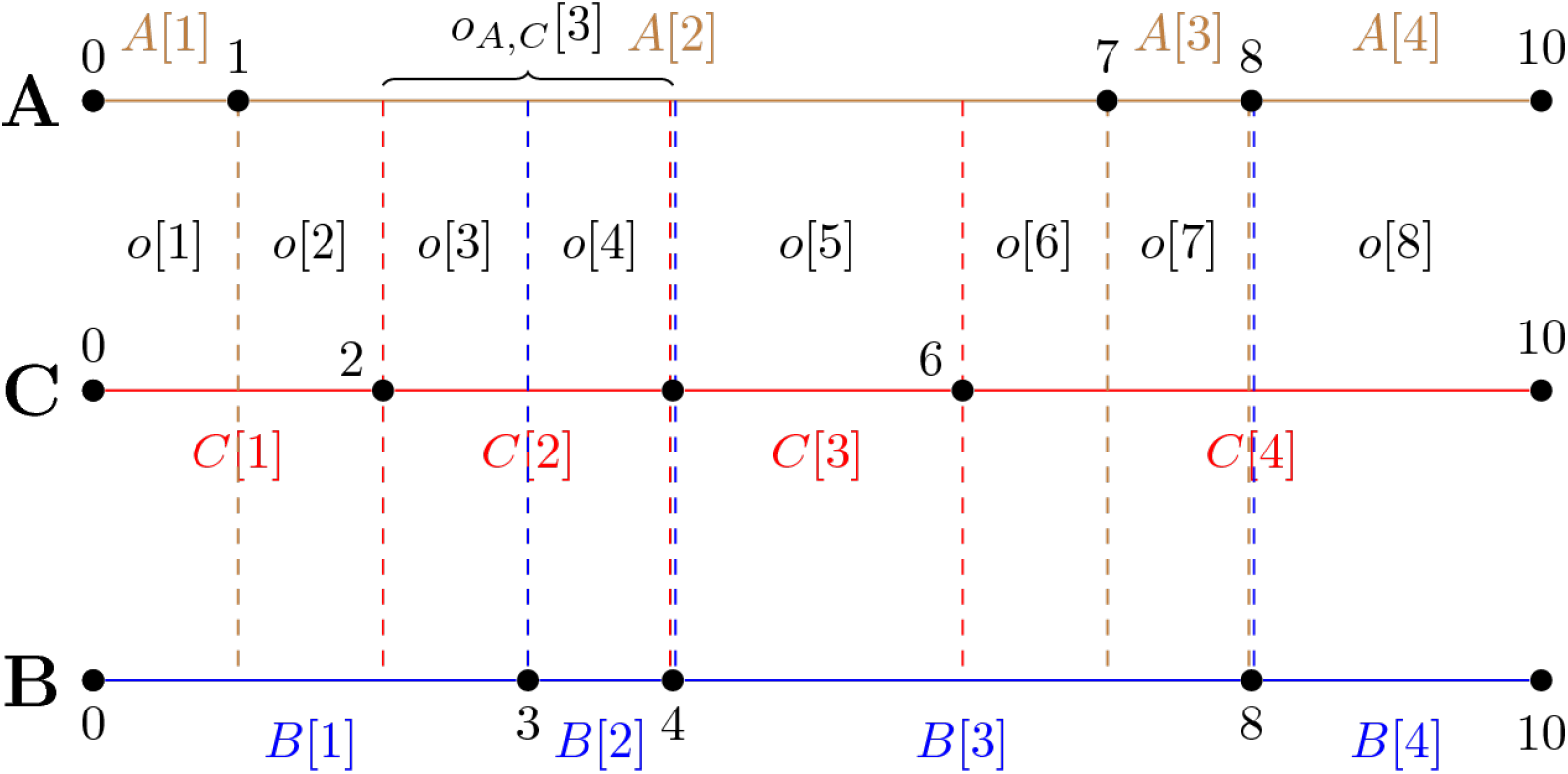
Ilustration of 3 partitions A, B, C and the induced segmentation.

**Definition 4.** *We define the function f*_*X,Y*_ (*i*), *which gives the original segment from partition X, that included the segment o*[*i*]. *More formally:*

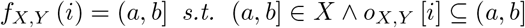

*We will use the same function to find segments in the second partition by writing f*_*Y,X*_.

**Definition 5** (BP distance). *Given 2 partitions X and Y s.t. l*(*X*) = *l*(*Y*) *their BP distance is defined as:*

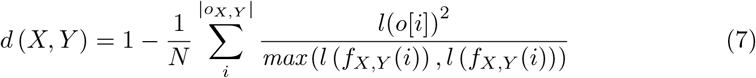

**Remark 3.** *From now on we will use A,B,C to distinguish between 3 partitions used to construct this proof and X,Y whenever we want to refer to any pair of partitions from set A,B,C s.t. X ≠ Y*.

**Claim 1.** *Function d satisfies: d*(*A, A*) = 0 *(identity of indiscernibles)*.

*Proof*.

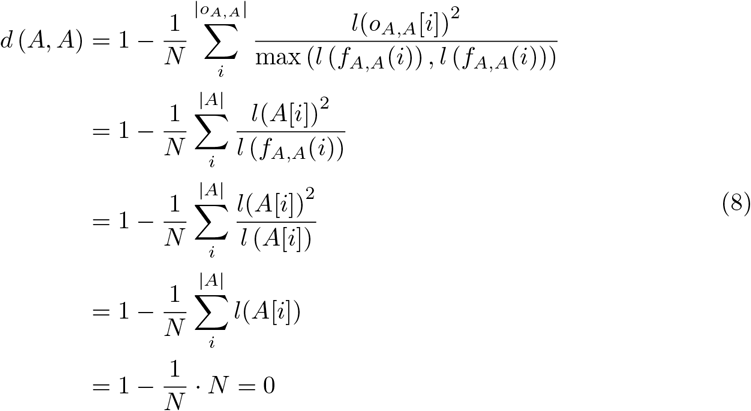

**Claim 2.** *Function d*(*X, Y*) *is symmetric for any A,B*

*Proof*.

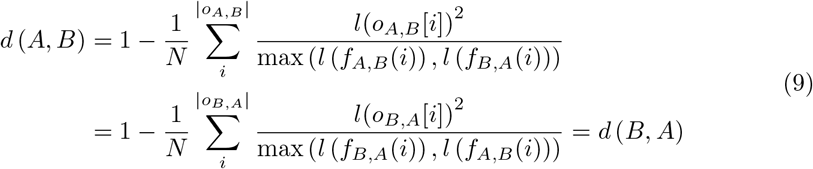

**Claim 3.** *Let us consider 3 domain sets A,B,C (refer to figure 6 as example). Function d satisfies:*

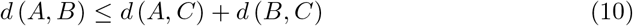

*Proof*. We start with putting equation 7 into inequality 10:

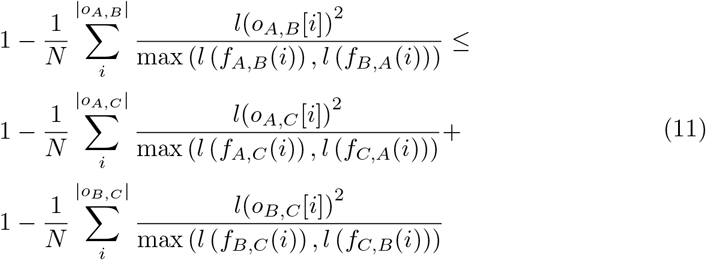

Using multinomial theorem we can substitute divisible segments with atomic ones in the following way:

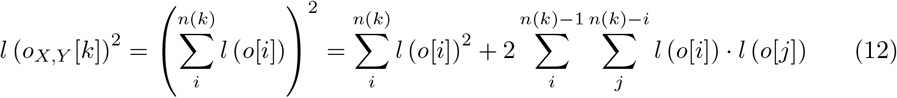

where: *n*(*k*) is the number of atomic segments in divisible segment *o*_*X*,*Y*_[*k*]. Obviously *n*(*k*) also depends on *X* and *Y*, but for simplicity we decided to leave it out of the notation here assuming it follows from the formula. Using equation 12 we can rewrite inequality 11:

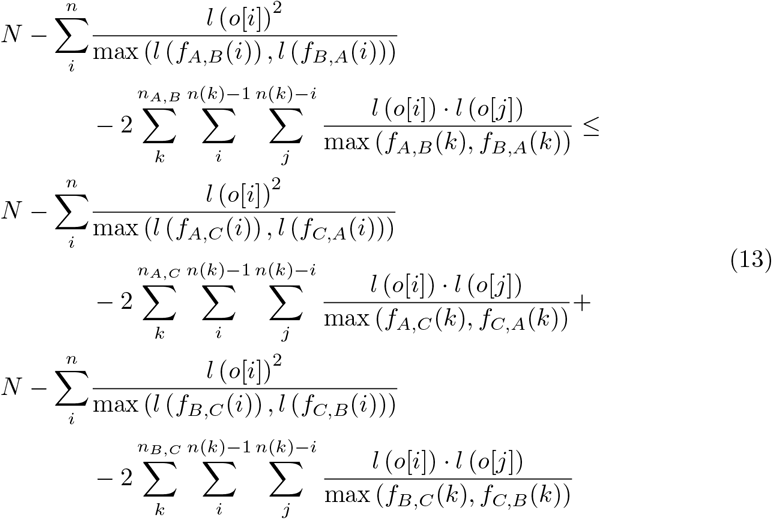

where: *n* is number of atomic segments and *n*_*X,Y*_ is number of divisible segments induced by *X*,*Y* partitioning. As atomic segments are common for *A, B* and *C* we can omit the subscript in *n*.

**Definition 6** (islands of segments). *We define a family of segments I*_*k*_ = {*o*[*i*]| *o*[*i*] ⊆ *C*[*k*]} *referred to as islands of segments, as satisfying the following condition:*

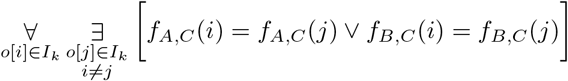

Intuitively islands of segments give rise to product sums on the right side of inequality 13.

**Claim 4.** *Any atomic segment o*[*i*] *satisfies:*

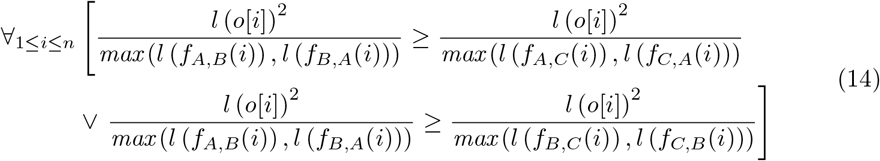

*Proof*.

1. first we can simplify notation by replacing *f_X,Y_* (*i*) with: *f_x_* (*i*) and *f*_*Y*,*X*_ (*i*) with: *f*_*Y*_ (*i*) as we are only concerned with atomic segments,
2. if *f_A_*(*i*) > *f_B_* (*i*), then:

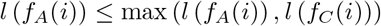
3. otherwise *f_A_*(*i*) ≤ *f_B_* (*i*) and:

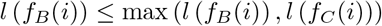

This allows us to split squared terms from the right hand side of inequality 13 and merge them into 2 groups (*S* - smaller, *R* - remaining), both of size *n*:

- *S* is sum of terms from either *A*,*C* or *B*,*C*, such that each term satisfies condition 2 (if it is a term from *A*,*C*) or condition 3 (if it is a term from *B*,*C*),
- *R* are remaining terms, i.e. they may not satisfy the above conditions.

We can rewrite inequality 13:

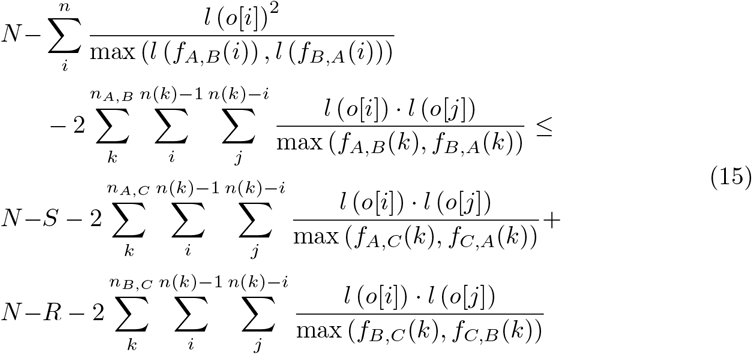

Now:

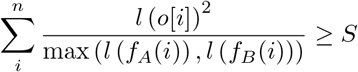

so we know that:

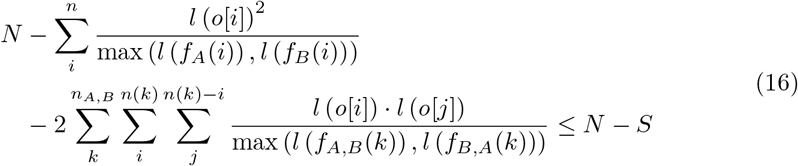

After using 16 to simplify 15, what is left to show is that:

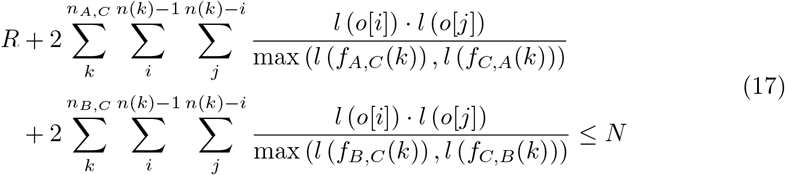

We note that no two segments *o_A,C_* [*k*] and *o*_*B*,*C*_ [*l*] can generate two different atomic segments *o*[*i*], *o*[*j*] that would be properly included in them.

**Claim 5.** *There are no 2 segments o_A,C_* [*k*] *and o_B,C_* [*l*], *such that for two different indexes i,j (i < j): o*[*i*] ⊂ *o*_*A,C*_[*k*] ∧ *o*[*j*] ⊂ *o*_*A,C*_[*k*] *and o*[*i*] ⊂ *o_B,C_*[*l*] ∧ *o*[*j*] ⊂ *o_B,C_*[*l*].

*Proof*.

1. assume that there are such 2 segments, which means that:

a. *s*(*o*_*A,C*_[*k*]) ≤ *s*(*o*[*i*]) < *e*(*o*[*i*]) ≤ *s*(*o*[*j*]) < *e*(*o*[*j*]) ≤ *e*(*o*_*A,C*_[*k*])
b. *s*(*o*_*B,C*_[*l*]) ≤ *s*(*o*[*i*]) < *e*(*o*[*i*]) ≤ *s*(*o*[*j*]) < *e*(*o*[*j*]) ≤ *e*(*o*_*B,C*_[*l*])
2. if *o*[*i*] ⊂ *o_A,C_* [*k*] ∧ *o*[*j*] ⊂ *o_A,C_*[*k*] then:

a. ∃*u*,*v*,*u*≠*v e*(*B*[*u*]) = *e*(*o*[*i*]) ∧ *s*(*B*[*v*]) = *s*(*o*[*j*])
3. but 2 contradicts 1b as by definition of *o*_*B,C*_[*l*] we have:

a. ∃_*w*,*z*_*s*(*o*_*B,C*_[*l*]) = max(*s*(*B*[*w*]), *s*(*C*[*z*]))
b. ∃_*w*,*z*_*e*(*o*_*B,C*_[*l*]) = min(*e*(*B*[*w*]), *e*(*C*[*z*]))

so: *s*(*B*[*w*]) < *e*(*o*[*i*]) and *s*(*o*[*j*]) < *e*(*B*[*w*])

Now we merge both product sums from inequality 17 and split them into *m* groups *P*_*k*_ corresponding to islands *I*_*k*_:

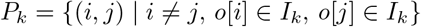

We also simplify notation:

1. as we consider islands of segments we can replace *f_C,A_*(*k*) and *f*_*C*,*B*_(*k*) with *f*_*C*_(*k*),
2. for any (*i,j*) ∈ *P*_*k*_ we introduce the notation *f*_*A*∨*B*_(*i*):

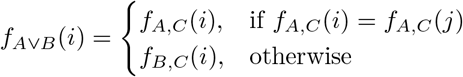

Rewriting inequality 17 gives:

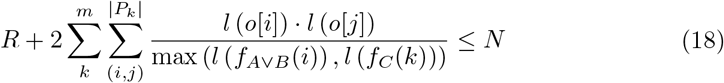

The number of elements in *P*_*k*_ can be upperbounded by:

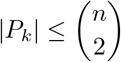

This allow us to upperbound the left hand side of inequality 18:

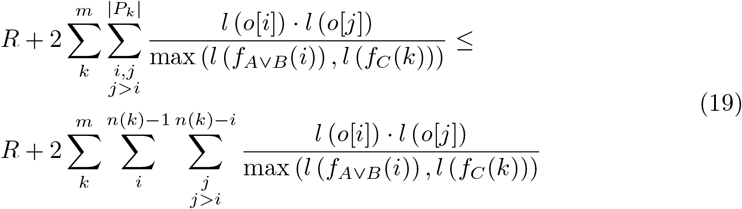

We can now split *R* on 2 groups:

1. segments *o*[*i*] such that: 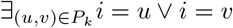,
2. remaining segments,

So we can write:

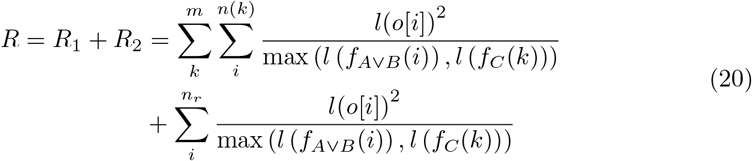

Now we put equation 20 into the right hand side of inequality 19:

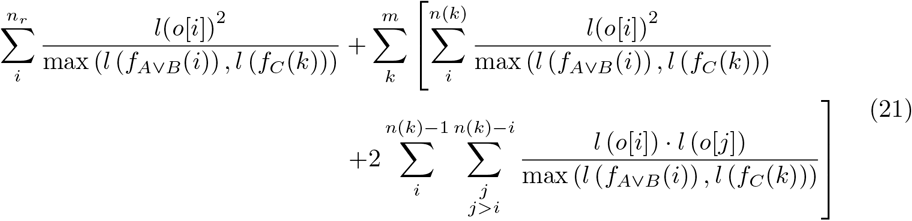

In order to further upperbound the left hand side of inequality 21 we need to select the minimum possible denominator. It can be easily shown that it is minimum when:

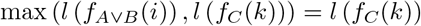

This follows from the definition of islands of segments as each atomic segment *o*[*i*] ∈ *I_k_* also satisfies: *o*[*i*] ⊂ *C*[*k*] meaning *l*(*f*_*A*∨*B*_(*i*)) ≤ *l*(*f*_*C*_(*k*)). The latter upperbound let us write again:

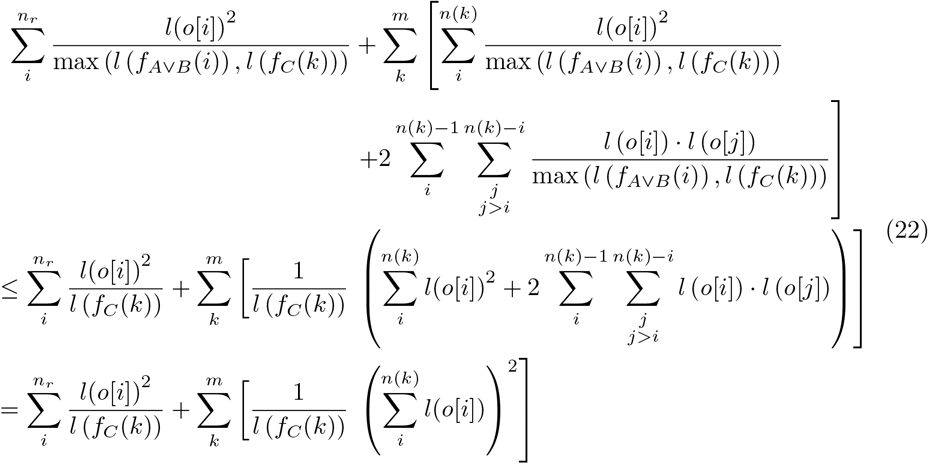

We can further upperbound 22 by selecting minimum *l*(*f_C_*(*k*)):

1. min(*l*(*f*_*C*_(*k*))) = *l*(*o*[*i*]) for atomic segments,
2. 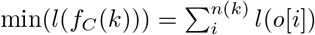 for divisible segments.

And we rewrite 22:

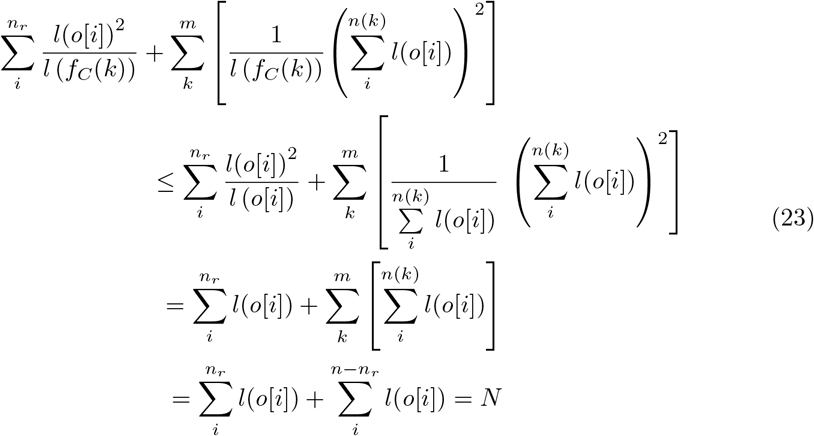

Which is what we wanted to prove. □

## Supporting information

**S1 Fig. Technical and biological replicates distance distribution of real Hi-C datasets by chromosome using different metrics.**

**S2 Fig. Comparison of p-values between technical and biological replicates distance distribution of different chromosomes using different metrics.**

**S3 Fig. Relationship between gene expression fold change and local BP score or local MI score of all genes for different pairs of cell types.**

**S4 Fig. Relationship between gene expression fold change and local BP score or local MI score of all genes assigned to 30 quantiles for different pairs of cell types.** Shown is mean gene expression fold change for each quantile.

**S5 Fig. Relationship between gene expression fold change and local BP score or local MI score of all genes assigned to 30 quantiles for different pairs of cell types.** Shown is median gene expression fold change vs median local score for each quantile as well as their Pearson corelation.

**S6 Fig. Corelation between median quantile gene expression fold change and median quantile local score for different pairs of cell types.**

**S7 Fig. P value of corelation between median quantile gene expression fold change and median quantile local score for different pairs of cell types.**

**S8 Fig. Relationship between methylation fold change and local BP score or local MI score of all TADs for different pairs of cell types.**

**S9 Fig. Relationship between methylation fold change and local BP score or local MI score of all TADs assigned to 30 quantiles for different pairs of cell types.** Shown is mean methylation fold change for each quantile.

**S10 Fig. Relationship between methylation fold change and local BP score or local MI score of all TADs assigned to 30 quantiles for different pairs of cell types.** Shown is median methylation fold change vs median local score for each quantile as well as their Pearson corelation.

**S11 Fig. Corelation between median quantile methylation fold change and median quantile local score for different pairs of cell types.**

**S12 Fig. P value of corelation between median quantile methylation fold change and median quantile local score for different pairs of cell types.**

**S1 Table. Corelation between median quantile gene expression fold change and median quantile local score for different pairs of cell types.**

**S2 Table. Corelation between median quantile methylation fold change and median quantile local score for different pairs of cell types.**

## Acknowledgments

This work was supported by the Polish National Science Centre grant decision No. [DEC 2015/16/W/NZ2/00314].

